# Modeling Hippocampal Spatial Cells in Rodents navigating in 3D environments

**DOI:** 10.1101/2022.06.30.498249

**Authors:** Azra Aziz, Bharat Kailas Patil, Kailash Lakshmikanth, Peesapati S S Sreeharsha, Ayan Mukhopadhay, V Srinivasa Chakravarthy

**Author notes:** **Correspondence:** V Srinivasa Chakravarthy. These authors contributed equally to this work (Bharat Kailas Patil, Kailash Lakshmikanth).

## Abstract

Studies on the neural correlates of navigation in 3D environments are plagued by several unresolved issues. For example, experimental studies show markedly different place cell responses in rats and bats, both navigating in 3D environments. In an effort to understand this divergence, we propose a deep autoencoder network to model the place cells and grid cells in a simulated agent navigating in a 3D environment. We also explore the possibility of a vital role that Head Direction (HD) tuning plays in determining the isotropic or anisotropic nature of the observed place fields in different species. The input layer to the autoencoder network model is the HD layer which encodes the agent’s HD in terms of azimuth (θ) and pitch angles (ϕ). The output of this layer is given as input to the Path Integration (PI) layer, which integrates velocity information into the phase of oscillating neural activity. The output of the PI layer is modulated and passed through a low pass filter to make it purely a function of space before passing it to an autoencoder. The bottleneck layer of the autoencoder model encodes the spatial cell like responses. Both grid cell and place cell like responses are observed. The proposed model is verified using two experimental studies with two 3D environments in each. This model paves the way for a holistic approach of using deep networks to model spatial cells in 3D navigation.

## Introduction

Spatial navigation is a competency that is crucial for an organism’s survival. A sizeable of body of literature that seeks to study neural substrates for spatial navigation focuses on the hippocampus, thanks to the popularity of this system as the “GPS of the brain” (Taube, Muller and Ranck, 1990; Burgess, Recce and O’Keefe, 1994; Hafting *et al*., 2005). Neuroscience research dedicated to spatial navigation and hippocampus has invested a far greater effort in 2D navigation than in 3D navigation perhaps to the practical constrains involved in the study of the latter (Bingman and Able, 2002; Geva-Sagiv *et al*., 2015). The recordings of hippocampal neurons that potentially encode space are frequently taken from surface-dwelling animals like rats (Hayman *et al*., 2011; Grieves *et al*., 2020, 2021) and only less often in flying creatures like bats (Yartsev, Witter and Ulanovsky, 2011; Yartsev and Ulanovsky, 2013; Eliav *et al*., 2021) to understand spatial maps in 3D.

Concepts that are relatively well established in case of 2D navigation run into rough weather when extended to the 3D case. For example, the basis of neural encoding of head direction (HD) in 3D space is still unclear, as experimental studies from different species yield conflicting reports. Shinder and Taube (Shinder and Taube, 2019) proposed that the HD cells of rats from the antero-dorsal thalamus are mainly tuned to the horizontal directions, and tuning is intact as long as the pitch angle is less than 90 degrees. On the contrary, Laurens and Angelaki (Laurens and Angelaki, 2019) countered the claim and proposed that the HD cells are tuned in all three dimensions of angular space. Finkelstein (Finkelstein *et al*., 2014) proposed that the HD is tuned to azimuth and pitch/roll or to the conjunctive combination of directions in bats. These discrepancies in results can be due to the distinctive navigational capabilities of rats and bats in their native environments (Barry and Doeller, 2013). Since bats fly, they map the environment volumetrically; hence, HD cells are tuned to combinations of azimuth and pitch or roll. Whereas since rats generally dwell on the ground, their HD cell tunings are predominantly limited to azimuthal angles and are less sensitively dependent on the pitch angle. Kim and Maguire (Kim and Maguire, 2019) demonstrated, using virtual reality (VR) experiments in humans, that the anterior thalamus and subiculum encode HD in the azimuthal plane while sensitivity for pitch directions is observed in the retro-splenial cortex.

The early discovery of place cells and grid cells in the hippocampus had inspired an extensive search for a comprehensive cellular system in the hippocampus for encoding space. Many experimental studies in bats have reported 3D place cells (Yartsev and Ulanovsky, 2013; Rubin, Yartsev and Ulanovsky, 2014; Grosmark and Buzsáki, 2016; Wohlgemuth, Yu and Moss, 2018). On the contrary, reports of grid cells in bats from the entorhinal cortex are relatively rare (Yartsev, Witter and Ulanovsky, 2011). Moreover, the grid cells in bats were recorded as the bat crawls on a 2D surface, and the gridness score was comparable with the grid cells of rats in a 2D environment. Therefore, there is a dearth of volumetric studies for grid cells in 3D as the aforementioned study is the only experimental study to record from such cells in bats.

There are experimental studies on rats in 3D environments, where place cell and grid cell mapping on the z-axis (gravity axis) is limited (Hayman *et al*., 2011; Grieves *et al*., 2020). (Grieves *et al*., 2020) showed that the place cells are significantly elongated along the gravity axis in an aligned lattice maze (aligned to the ground). While in the case of a tilted lattice maze, the place fields were more isotropic as all three axes were topographically equivalent. The rationale provided for the observation was that for the aligned lattice, the animal was not moving freely in the gravity axis, whereas the motion was equiprobable for all axes in the tilted lattice. Similarly, Hayman and colleagues (Hayman *et al*., 2011) reported elongation of place fields and grid fields on the gravity axis in pegboard and helical mazes.

In this study, we focus on the following set of questions in 3D navigation:

- We propose a versatile deep autoencoder based network to model different spatial cells observed in hippocampal formation in diverse 3D environments.
- We demonstrate the dependence of isotropic and anisotropic nature of spatial cells on HD tuning for different species of animals.

Although there are several computational models of hippocampal spatial cells in relation to 2D environments (Solstad, Moser and Einevoll, 2006; Hasselmo, Giocomo and Zilli, 2007; Burak and Fiete, 2009; Bush and Burgess, 2014; Soman, Muralidharan and Chakravarthy, 2018), very few models are available for 3D navigation. Mathis et al (Mathis, Stemmier and Herz, 2015) described a mathematical model to show that the Face Centered Cubic (FCC) lattice is the most optimal representation for equivalent grid cells in a 3D environment. A computational study (Stella and Trevesy, 2015) suggests the possibility of the formation of grid cells in FCC or Hexagonal Closed Packing (HCP) type of lattice configuration. Since the oscillatory interference models have intrinsic theta oscillations, and theta rhythms are not observed in freely flying bats, Horiuchi and Moss (Horiuchi and Moss, 2015a) proposed a four-ring attractor network. The network uses the interference of spatial patterns from the ring attractors to construct grid patterns in 3D, assuming a reference vector separated by 109 degrees. Soman et al (Soman, Chakravarthy and Yartsev, 2018) proposed an anti-Hebbian network to model place cells in freely flying bats and studied the effect of the range of pitch angles on the isotropic nature of place cells and grid cells. The model’s neurons also imitated border cells and theorized the presence of *plane cells*.

Deep learning is a widely accepted technique to solve a plethora of problems in diverse fields, including computer vision (Voulodimos *et al*., 2018), natural language processing (Young *et al*., 2018), reinforcement learning (Arulkumaran *et al*., 2017), etc. It has successfully proven its ascendency for object recognition tasks (Rashid *et al*., 2020). For a long time, the relevance of deep network models like the Convolutional Neural Networks (CNNs) for neuroscience research has been thought to be limited to learn input output behavior. However, recent studies have shown that even the activation patterns of the layers of CNNs can be gainfully compared to the activation patterns of cortical areas in primary visual and auditory systems (Kell *et al*., 2018). These modeling studies have inspired application of deep learning networks to model spatial cells in 2D and related navigational problems (Banino *et al*., 2018; Cueva and Wei, 2018; Aziz *et al*., 2021). However, there is a paucity of studies that apply deep learning techniques to model hippocampal responses in 3D.

We propose an autoencoder-based deep network to study spatial cells in rats navigating through a 3D environment. The path integration (PI) process in our model is inspired by (Soman, Muralidharan and Chakravarthy, 2018), but the time dependency of neural responses in the model is discarded by an averaging process. The high-frequency components from the PI neurons in the model are removed using a low pass filter to make it purely a function of space. The grid cell and place cell responses emerge in the hidden layer of the model. Amongst all the available models of spatial cells in 3D environments, none have attempted to design a comprehensive model to account for a variety of experimental studies in 3D navigation. In this paper, we primarily model two distinctive experimental studies (Hayman *et al*., 2011; Grieves *et al*., 2020), each having two setups, an aligned and tilted lattice maze and a pegboard and helical maze.

The paper is outlined as follows. The following Methods section, starts with data generation in terms of trajectories used in different environments. Then the model architecture and the training procedure are described. The subsequent section describes the simulated results from two experimental studies (Hayman *et al*., 2011; Grieves *et al*., 2020) and their statistical inferences. Lastly, we summarize the study in the discussion section.

## Methods

The aim of the proposed study is to model the 3D spatial navigation experiments of (Hayman *et al*., 2011; Grieves *et al*., 2020). These experimental studies use lattices of different geometries which are described below.

### Trajectory generation for 3D lattice maze

To simulate the study of (Grieves et al. 2020) which uses a cubical lattice maze, we use a lattice maze of an outer boundary size of 5×5×5 units. Furthermore, navigation experiments were conducted under two conditions of the lattice: aligned lattice and tilted lattice (Fig. 1). Each axis has six bars and the bars intersect to form nodes. The simulated agent’s movement from one node to the other is based on probabilities similar to experimental conditions (Grieves *et al*., 2020) where the rat moved predominantly parallel to the X or Y axis (ground-axes) and moved less in the vertical dimension (see Supplementary material: section 1) in case of aligned lattice maze (Fig. 1: A). When the agent moves from one node to the other, it mostly moves within its 180 degrees horizontal field of view. The agent moves in the vertical direction (upwards or downwards) with a probability of 20% and on the horizontal axis with a probability of 80%. Whereas in a tilted lattice maze, the agent moves with equal probabilities on all the three axes (Fig. 1: B, see Supplementary material: section 1). The resulting sequence of nodes was smoothened using a cubic spline to form a smooth trajectory. Speed is assumed constant throughout and T number of equidistant points are generated along the trajectory.

**Figure 1:**
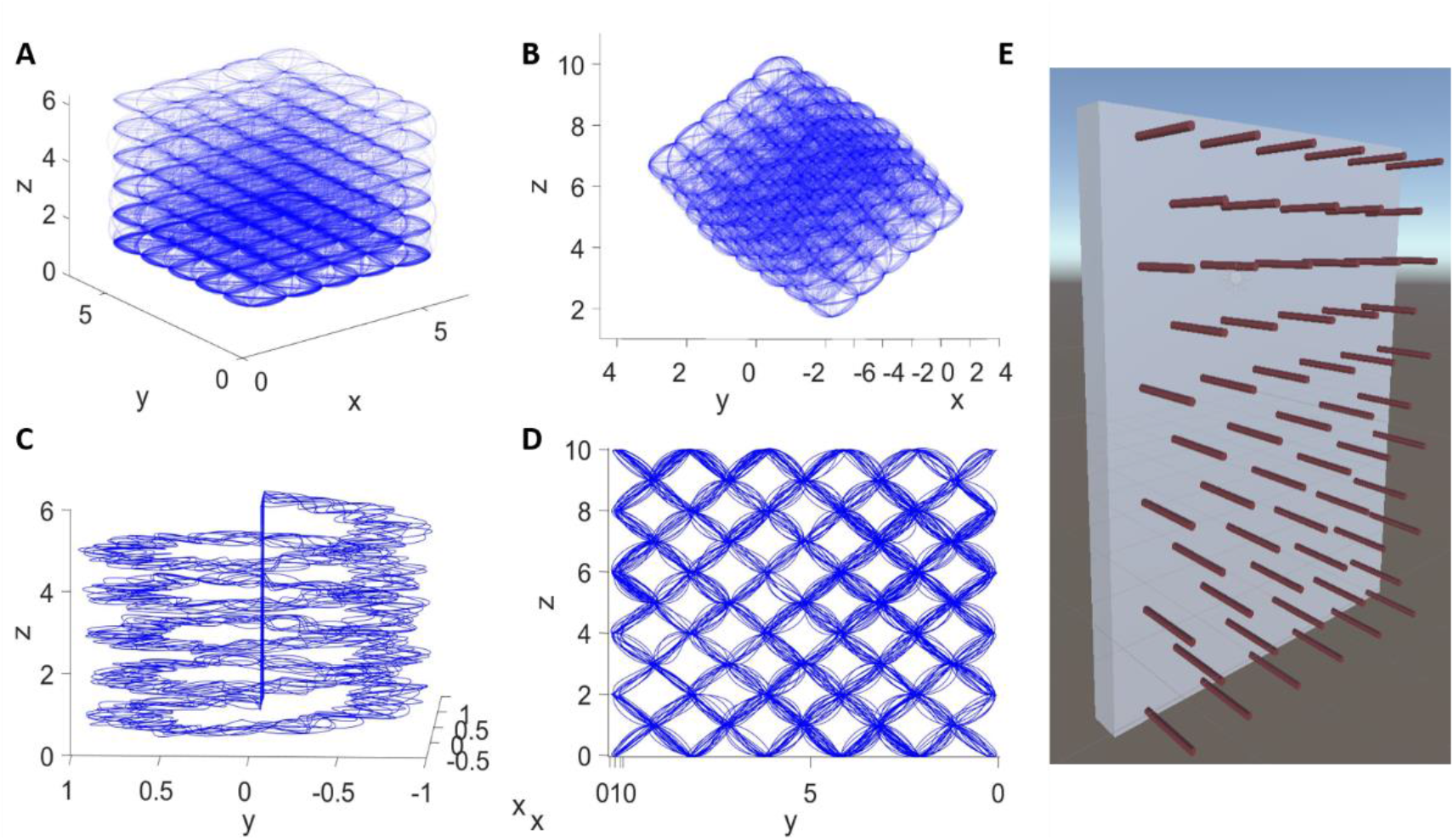
The virtual agent’s trajectory in A) aligned lattice maze B) tilted lattice maze. C) helical maze. D) pegboard maze. E) depicts the pegboard environment redrawn based on (Hayman et al., 2011).

### Trajectory generation for helical maze

The helical trajectory is generated with an assumption of a virtual agent similar to the lattice maze. This trajectory consists of five coils starting from the ground and extending to a height of five units increasing uniformly throughout the helix. The inner radius of the path for each coil is 0.5 units and the outer radius is 1 unit. In this path, the animal is allowed to move freely in any random trajectory within the above bounds (Fig. 1: C). These specific dimensions are employed to match the experimental conditions (Hayman *et al*., 2011). Moreover, the upward and downward trajectories are generated separately. For upward motion, the agent directly returns to the ground after reaching the top without any downward movement along the helix, and vice versa for the downward motion.

### Trajectory generation for pegboard maze

A trajectory on a pegboard of dimensions 1×10×10 units is created. Here the length of the peg is 1 unit whereas the pegboard is 10×10 units (Fig. 1: D and E). The agent moves with equal probability from any given node considering that it will not go back to the node it just came from. The agent jumps over different nodes taking 2,000 steps. Then the trajectory is interpolated to produce 50,000 points and smoothed using spline functions.

### Model Description

#### Head Direction (HD) layer

HD matrix represents the preferred directions of neurons in the azimuth (*θ*) and pitch (*ϕ*) plane. Here we consider n_1_ number of equally spaced azimuth angles spanning 360 degrees and n_2_ number of equally spaced pitch angles spanning 180 degrees. Using these angles, (n_1_×n_2_) unit vectors were used as the preferred directions of the HD neurons. The matrix of preferred directions is calculated as follows.

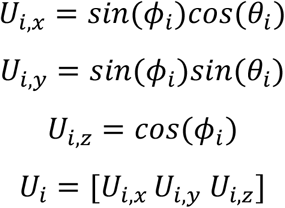

where, *U*_*i,x*_,*U*_*i,y*_,*U*_*i,z*_ are the components of HD in x, y and z axes for *i*^*th*^ direction. The response of the *i*^th^ HD cell, HD_*i*_, is given as

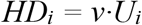

where *v* is the velocity of the agent in 3D space and *U*_*i*_ is the preferred direction.

#### PI layer

In the PI layer, the neural responses are functions of space and time (Soman, Muralidharan and Chakravarthy, 2018). As the spatial cell responses are typically depicted in terms of space alone, we remove the time dependency from the PI neurons by an averaging process as discussed in (Aziz et al, 2021). The neurons in the PI and the HD layers have a one-to-one connection. The response of the i^th^ neuron of the PI layer is given as,

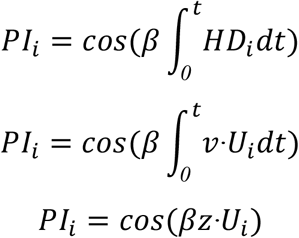

where *z* is the displacement of the agent from the initial position and *β* is the modulation factor. The output of the PI layer is passed on to the autoencoder layer.

#### Autoencoder Model

The autoencoder is an unsupervised neural network model used to perform dimensionality reduction (Wang, Yao and Zhao, 2016). The output from the PI layer is passed on to the autoencoder network through feedforward connections. We consider two variations of the autoencoder network: one with a single layer (Fig. 2: A) and one with two layers (Bengio *et al*., 2006; Aurelio *et al*., 2007; Wang, Yao and Zhao, 2016)(Fig. 2: B). The hyperparameters used for training the autoencoder network are shown in Table 1 for all experiments.

**Figure 2:**
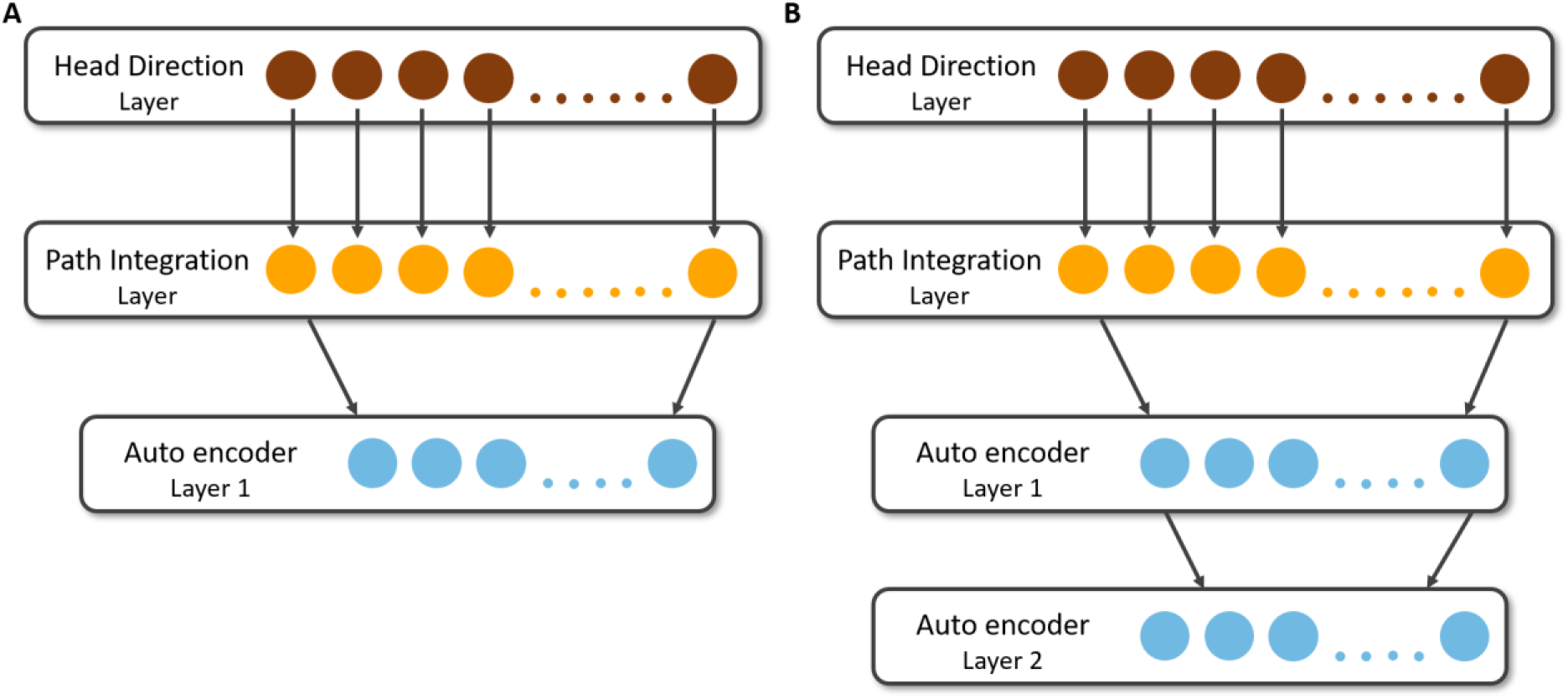
A schematic of the Autoencoder Model. Circles represent neurons. A) Architecture of the model with single autoencoder layer. B) Architecture of the stack autoencoder model with two layers.

**Table 1:** Parameters and Hyperparameters used in the Autoencoder Network.

| Parameter | Value |
| --- | --- |
| $\beta$ (lattice maze) | 1.8 |
| $\beta$ (helical maze) | 4 |
| $\beta$ (pegboard maze) | 1.6 |
| $\Theta$ (azimuth angle) | $0-2\pi$ |
| $\Phi$ (pitch angle) | $0-\pi$ |
| $n_1$ (no. of preferred azimuth angles) | 20 |
| $n_2$ (no. of preferred pitch angles) | 3 |
| $n$ (hidden neurons in autoencoder model) | 50 neurons |
| Activation function | Linear |

## Results

### Simulation Studies

The common underlying theme of the simulation studies described below is to let the agent run through the lattice following a predetermined trajectory, train the autoencoder network described in Method Section, and examine the responses of the neurons in the autoencoder layer(s) for space encoding properties.

#### 3D lattice maze

The simulations are done for two lattice mazes: aligned lattice and tilted lattice maze. The results are discussed simultaneously for both cases. For this study, we use a single layer autoencoder (Fig. 1: B). The output of the hidden layer with 50 neurons of the autoencoder is saved for analysis to understand the place cell responses in a volumetric environment. The raw output of the neurons is thresholded suitably. The neurons which show non-place field-like firings are screened manually by visual inspection of their firing fields. Trajectory points where the activity of a neuron crosses the threshold are indicated by red dots (Fig. 3: A1 and A2). In the case of the aligned lattice maze, out of the 50 hidden neurons, 44% of neurons show responses similar to place cells. Whereas, in a tilted lattice maze, out of the 50 hidden neurons, 100% of the neurons show a place cell-like response. These place cells are further analyzed using firing rate maps.

**Figure 3:**
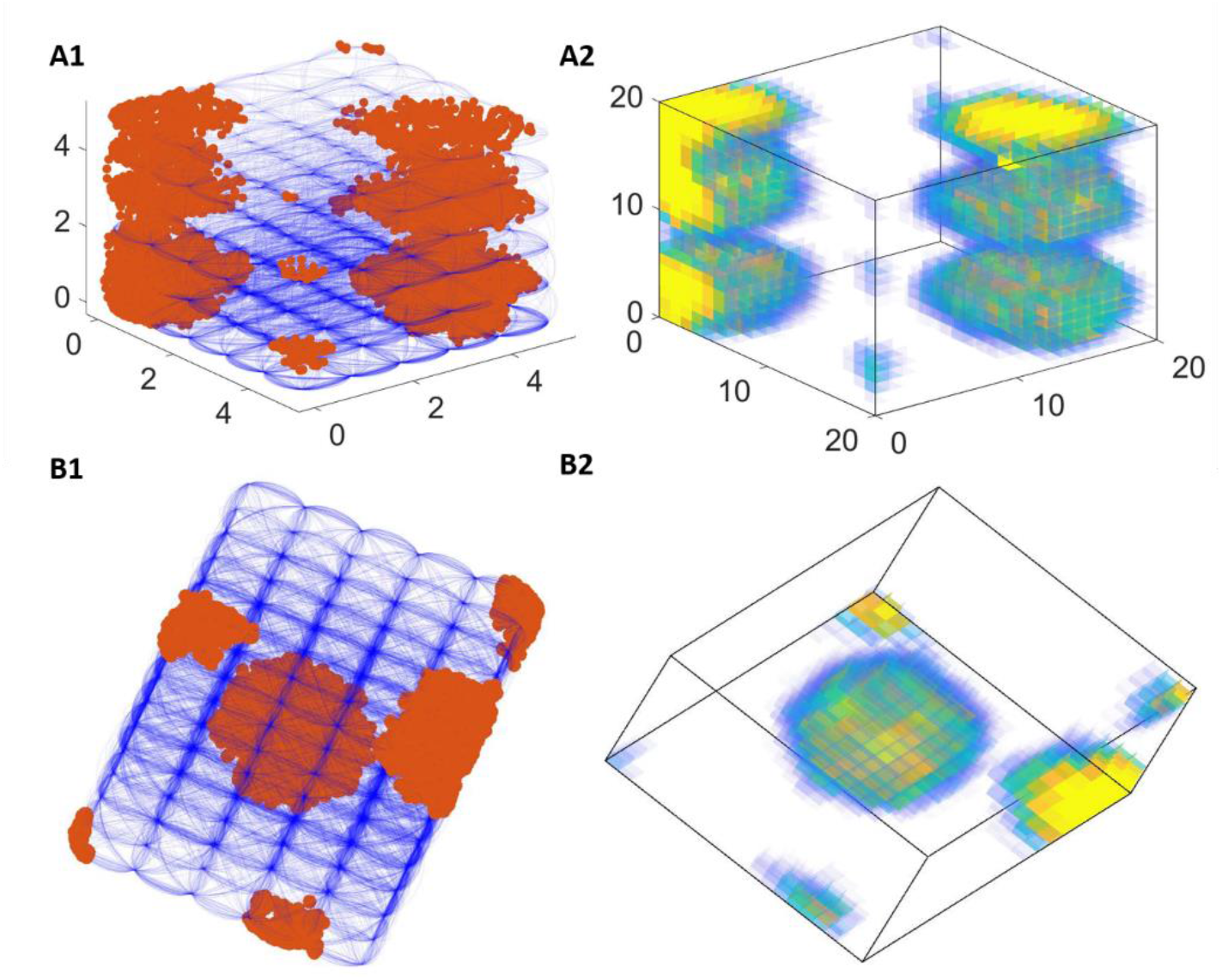
Firing field (A1) and firing rate map (A2) of a place cell in the aligned lattice maze. Firing field (B1) and firing rate map (B2) of a neuron in a tilted lattice maze.

To generate firing rate maps, the 5×5×5 unit^3^ environment is divided into 20×20×20 voxels. The firing activity is calculated using the thresholded data as specified in the experimental study (Grieves *et al*., 2020). The firing rate at the j^th^ voxel position *x*_*j*_ is defined as,

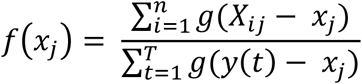

where, *X*_*ij*_ is the *i*^*th*^ firing point location in the firing field in *j*^*th*^ voxel, n is the total number of firing locations inside a voxel, *y*(*t*) is the position of the rat at a discrete-time step *t*, T is the total number of trajectory points, *x*_*j*_ is the position of the voxel center whose firing rate is being calculated. The truncated Gaussian function *g* gives non-zero values for neighboring 26 voxels.

Connected voxels are isolated by a MATLAB function “regionprops3” and are interpreted as place fields if the region is larger than 50 voxels (Fig. 3: A2 and B2). Some place cells are observed to have multiple place fields (Fig. 3: A2 and B2) identical to the experimental study (Grieves *et al*., 2020). The properties of place fields for both aligned and titled lattice mazes are then screened for further analysis.

#### Place fields were uniformly distributed in both mazes

To check whether the firing fields are uniformly distributed, we reproduced the distribution results observed in the experimental analysis. We observed that the median field centroids, generated by the autoencoder, lie close to the maze centers along each axis (Fig. 4: A1 and B1). The median of centroids along each axis lies within the 95% confidence intervals of a shuffle distribution generated by taking N uniform random points (where N is the number of place fields screened) within the lattice frame and calculating their centroids (Fig. 4 and Supplementary material: section 3).

**Figure. 4:**
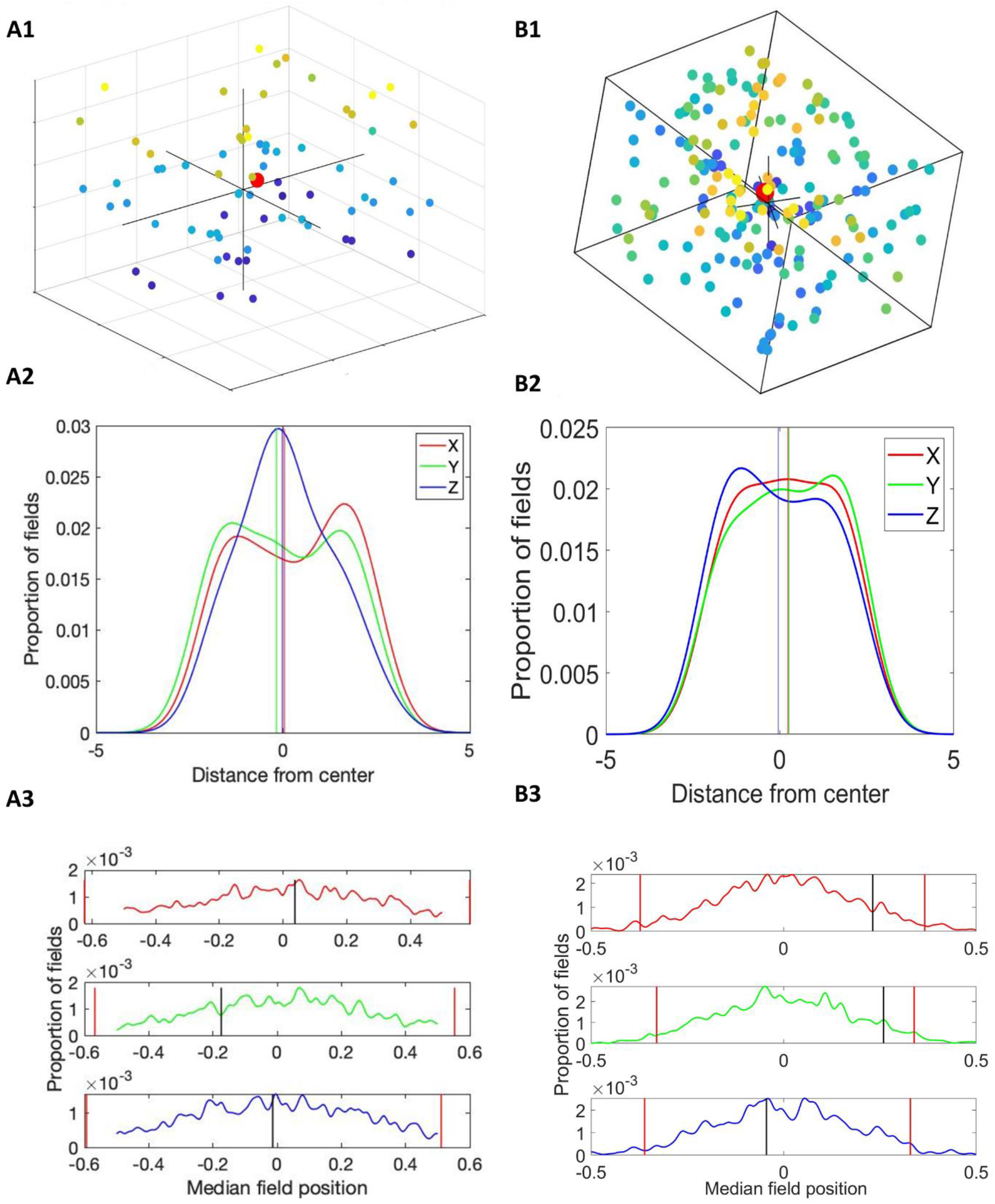
A) represents the aligned lattice. A1) Centroids of the place fields are represented by colored dots (hotness of plotted dots are directly proportional to the height in the maze. The red dot is the median of centroids of all place fields). A2) Distribution of place fields along each axis inside the volume. Lines represent the median for each axis. A3) Distribution created by generating N random points (equal to place field) and calculating their median for 1000 times. Red lines depict the 2.5th and 97.5th percentiles. Black lines represent the actual median of place fields. B) represents the same analysis for the tilted lattice case.

#### Place fields were elongated

Most firing fields were elongated in both the lattice mazes. Most elongation indices (see Supplementary material: section 4) deviated from 1 (elongation of 1 indicates a spherical place field). A small percentage (Fig. 5: A1 and B1 text) of place fields were more spherical than would be expected by chance in both cases (Fig. 5: A1 and B1).

**Figure 5:**
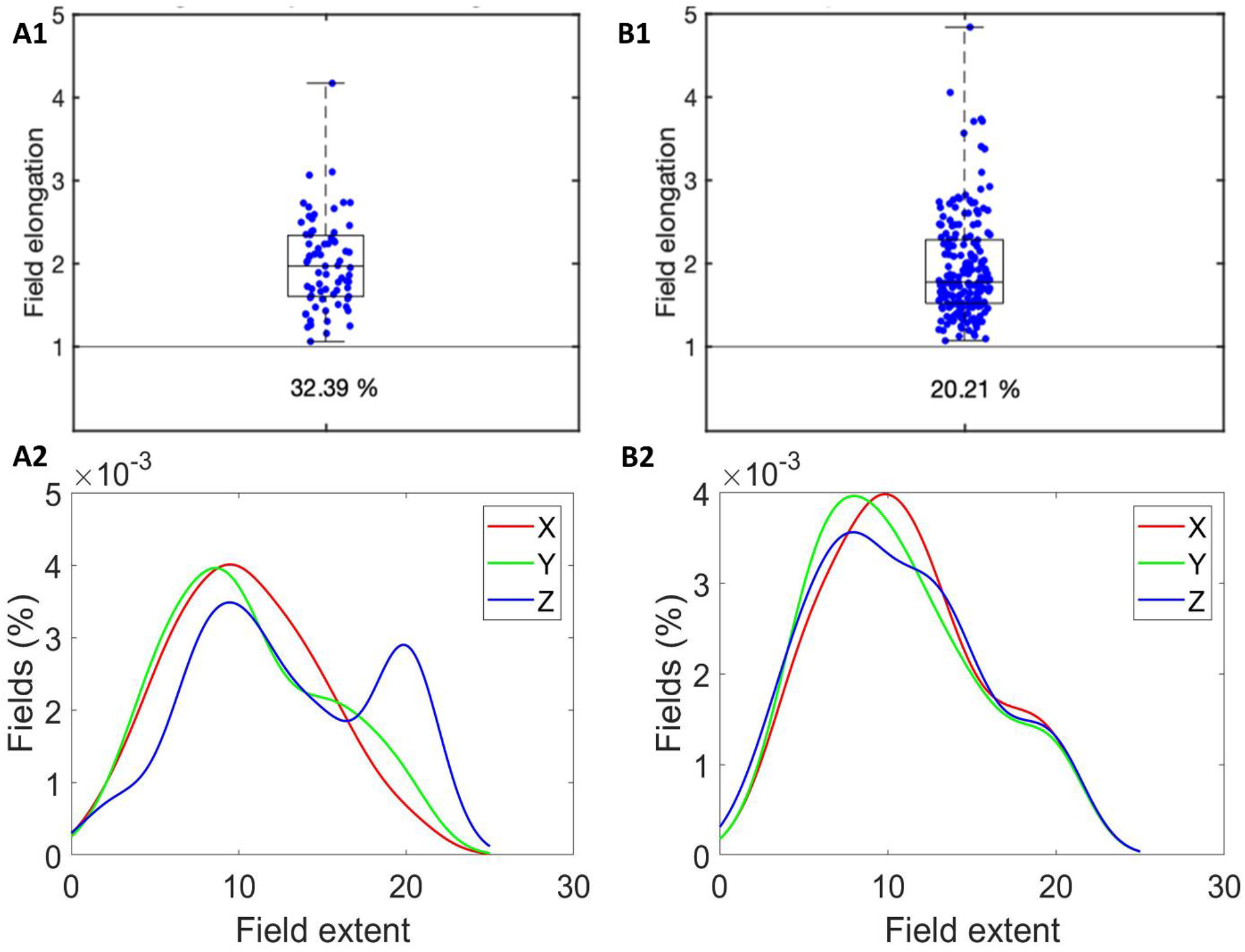
A) For the aligned lattice. 1) The elongation indices of all place fields are represented by blue dots. Index value of 1 (gray line in box plot) indicates a spherical field. 2) Distribution for the proportion of cells for field length along the respective axis. Z-axis in the aligned lattice has a significantly different bimodal occurrence. All the other axes have similar distributions. B) represents the same analysis for the tilted lattice.

The distribution of the proportion of fields showing the length of fields shared the same unimodal appearance across all axes in tilted and aligned except for Z-axis in case of the aligned lattice. Z-axis had bimodal occurrence with a second maximum for longer fields. This showed a significant elongation along the Z-axis compared to the others (Fig. 5: A2 and B2).

#### Place fields were elongated parallel to the maze axes

As in the last section, it is shown that the place fields are elongated in nature. But in order to understand the orientation of the elongated place fields, the orientation of the principal axis of the field is calculated and projected onto a unit sphere. In the aligned lattice maze, the majority of fields are oriented along the maze cardinal axes, although more fields are oriented along the Z-axis (Fig. 6: A1 and A3). In a tilted lattice maze, the majority of fields are oriented along tilted axes i.e. A, B, and C axes (Fig. 6: B1). (Fig. 6: A3 and B3) shows orientation of firing fields along specific axes. In aligned case, the firing field orientations only along X and Z axes are more than expected by chance (97.5^th^ percentile) (Fig. 6: A3) as compared to original experiment (Grieves *et al*., 2020) where all three axes (X, Y and Z) have firing field orientations expected more than by chance. In contrast, for the tilted lattice, the firing field orientations along all three tilted axes (A, B, and C) were observed more than expected by chance (Fig. 6: B3). In aligned case, XY and ABC axes are considered not different as they fall into the range of each other’s error bars (supplementary methods: section 5 - uniform sampling). But the X, Y, and Z axes compared to A, B, and C axes are significantly different in terms of field orientations for tilted lattice. To compare the field orientation between cartesian system and rotated system we calculated the ratio of field orientation (Fig. 6: A2 and B2) along both the systems. The observed results were found to be more than expected by chance (99^th^ percentile) in case of aligned lattice and less than expected by chance (1^st^ percentile) for titled lattice (Fig. 6: A2 and B2) (Supplementary material: section 5). This shows that the firing fields are more oriented towards the cartesian system in the aligned lattice and towards the rotated system in the tilted lattice.

**Figure 6:**
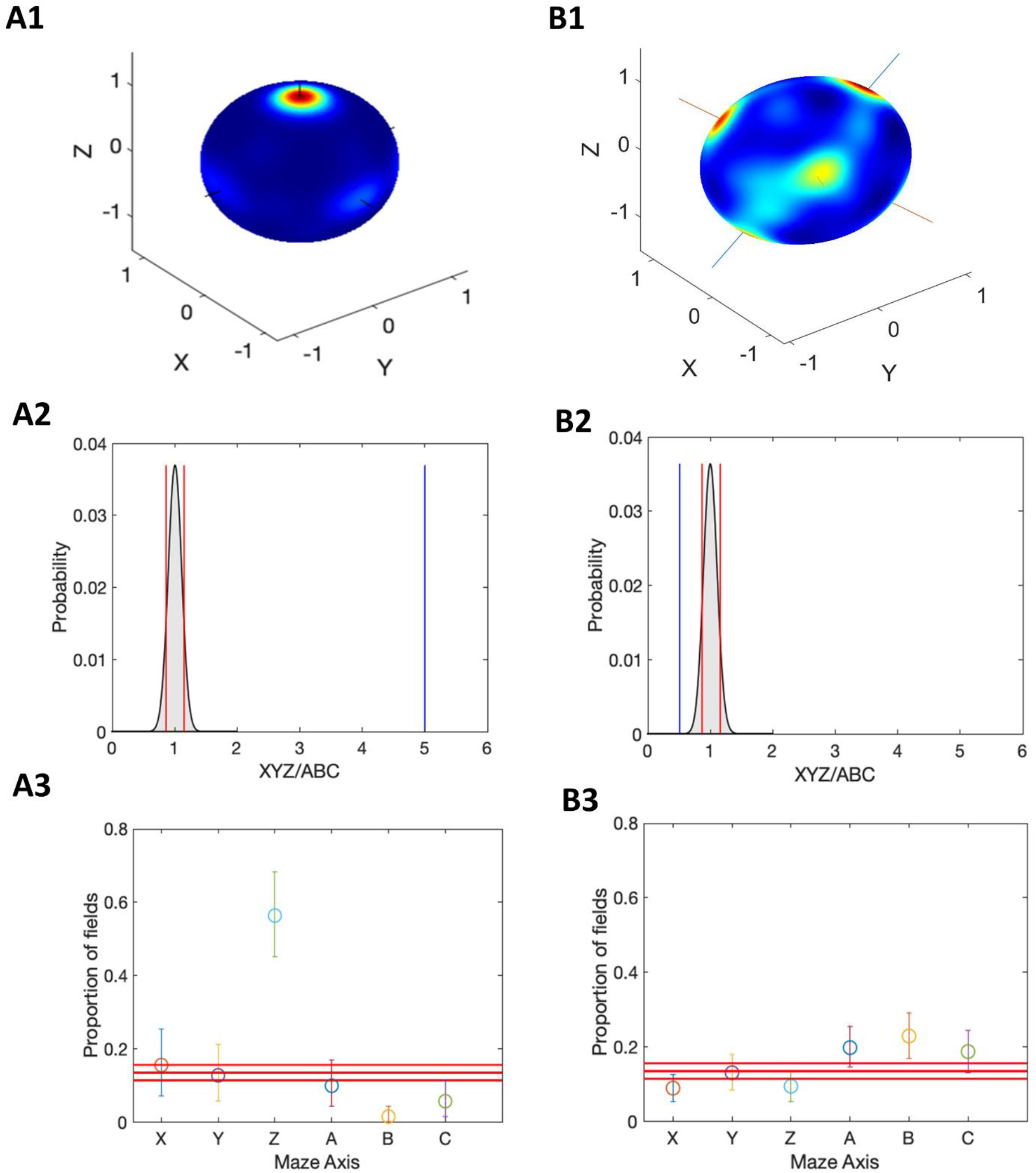
The place fields were oriented parallel to the maze axes. Hot colors represent the direction of the first principal axis of the place fields. A1) Place field orientation plot for aligned lattice maze. B1) Place field orientation plot for the tilted lattice maze. A2) distribution of the ratio of two axes system using shuffle test. Red lines represent 1^st^ and 99^th^ percentiles, blue line represents the actual value for aligned lattice. A3) Axis-wise orientation of the firing fields with 2.5^th^ and 97.5^th^ percentiles as error bars calculated using uniform sampling. The three red bars represent the 2.5^th^, 50^th^ and 97.5^th^ percentile of the firing fields expected by chance along any axis. B2, B3) represent the same analysis for the tilted lattice.

#### Spatial information along the axes

The aforementioned results show that the firing fields are elongated along the maze axes in both mazes. However, it is important to know if there is a preferential growth along any particular axis in both lattice mazes. It is observed that more place fields are elongated along the vertical (Z) axis in the aligned lattice maze, whereas there is no significant preference seen in the tilted lattice maze i.e, nearly equal fraction of firing fields are elongated along all the three axes (Fig. 7: A and B). Hence, the spatial information conveyed by the place cells in an aligned lattice maze is less along the Z-axis in aligned lattice maze. Using the binary morphology technique (supplementary material: section 6), we showed that vertical spatial coding was indeed less accurate than along the horizontal axes of the maze in the aligned lattice.

**Figure 7:**
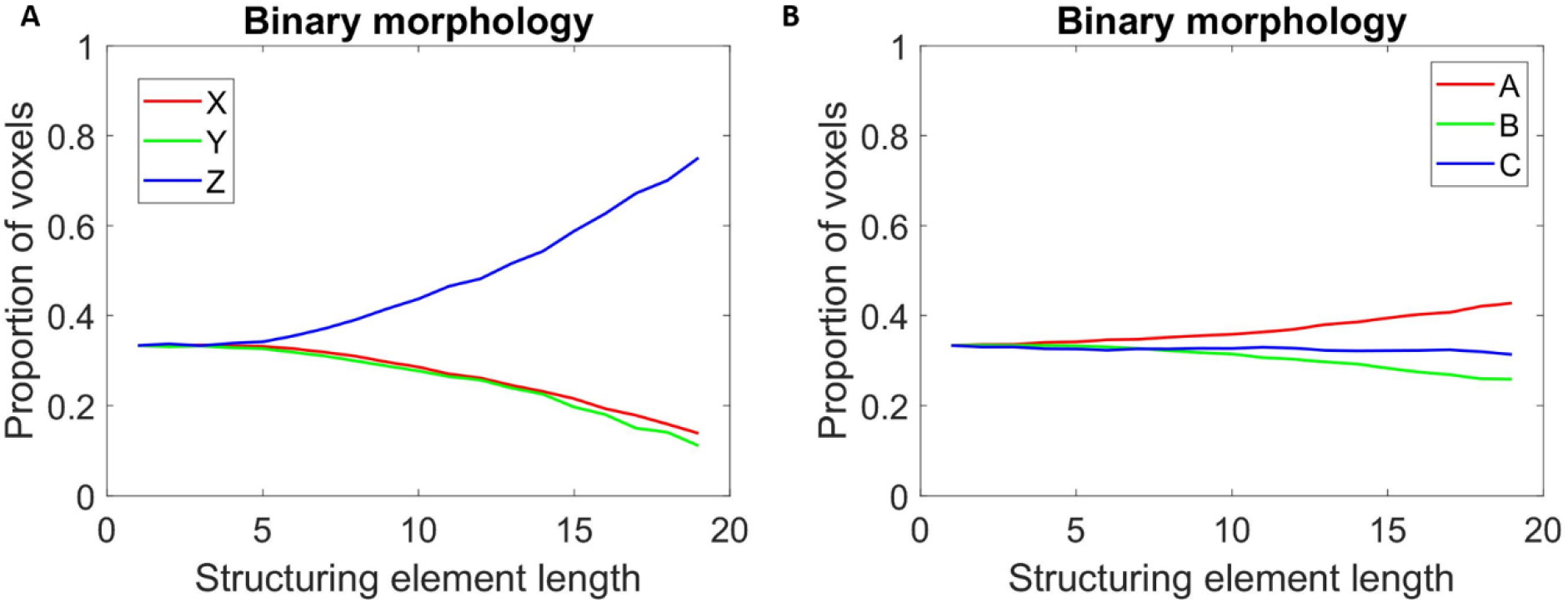
Binary morphological analysis of place cells’ firing rate maps. A) Initially, for smaller structuring element lengths, the connectivity is the same and then begins to diverge after erosion using structuring elements with lengths greater than 7. Hence, the voxels are highly connected along the Z-axis compared to the other cardinal axes, thereby suggesting a lower vertical spatial coding by place cells for the aligned lattice case. B) Axes in tilted lattice show equivalent proportion of field elongated along all.

#### Impact of HD Tuning

HD neurons are tuned for a set of azimuths (*θ*) and pitch (*ϕ*) angles. The azimuth is a uniform set of angles in the horizontal plane. The pitch is a set of angles with the axis parallel to the gravity axis. The preceding results are observed when the set of pitch angles is restricted only to the directions in which an agent can move. We expand this set by increasing the neurons that are tuned for different pitch angles in HD layer.

This expansion of pitch angle set, showed no significant impact on the distribution of place fields as the median of centroids in both (aligned and tilted lattice) stayed closer to the actual centroid of the mazes for all cardinal axes (Fig. 8: A1 and A2). The same observation extended to elongation indices too, as the expansion of pitch angle set showed no significant change in the distribution of elongation index for place fields (Fig. 8: B1 and B2).

**Figure 8:**
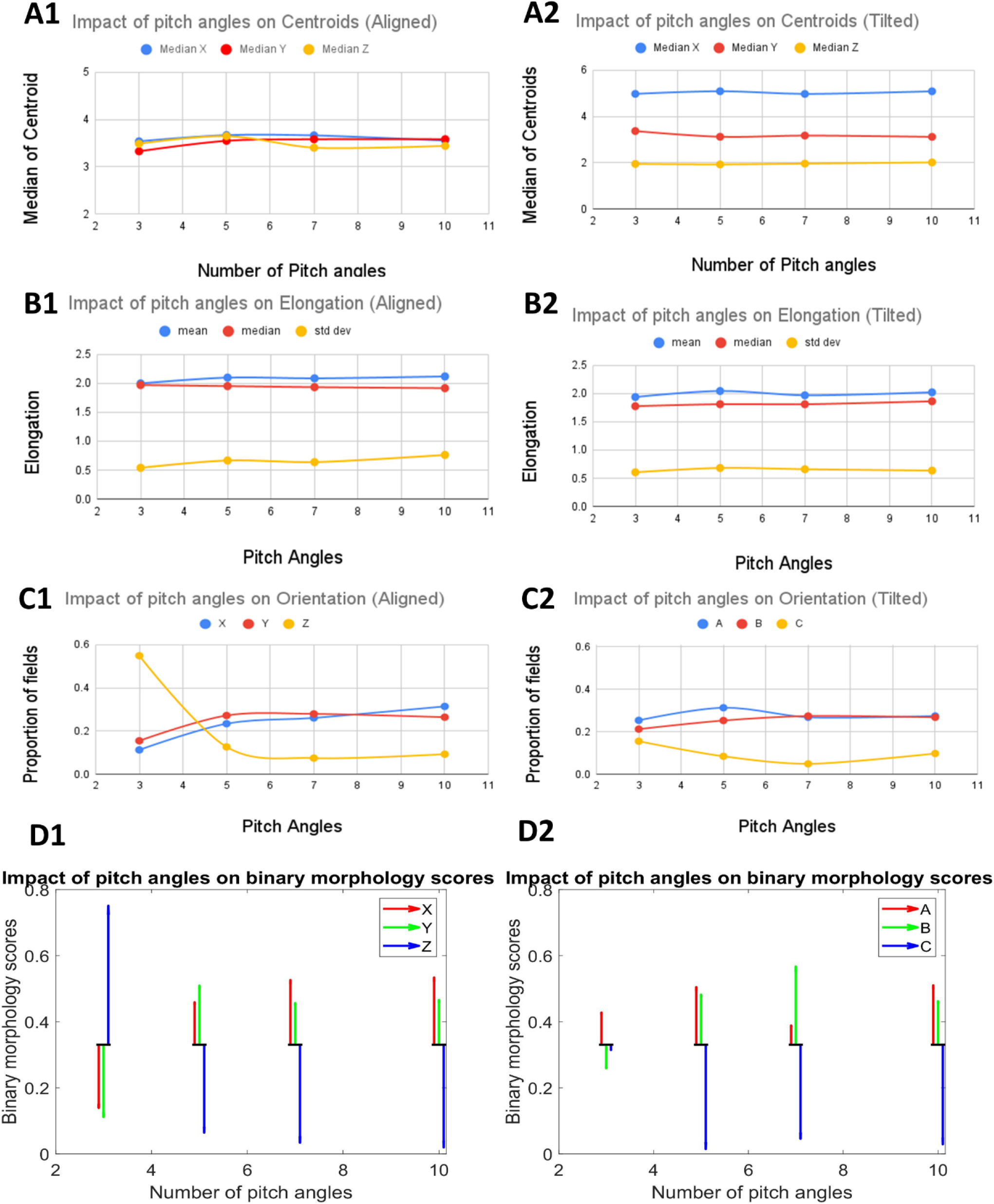
Impact of pitch angles on all phenomena observed in place fields. Left column, A1) Impact of HD tuning on Centroids of place fields for aligned lattice. B1) Impact of pitch angles on Elongation Index. Increase in number of pitch angles shows no change in both A1 and B1). C1) Impact of pitch angles on orientation of place fields along specific axes. Significant change is observed in orientation along Z axis. D1) Impact of pitch angles on binary morphology scores. Black lines denote the proportion of fields for the first structuring element. The direction of arrows denotes a general trend of proportions for increasing size of structuring elements. The length of the arrows signifies the range of increase/decrease in the proportion of fields along that specific axis.

An enormous proportion of place fields were elongated along Z axis when fewer pitch angles for HD neurons are employed. But this ratio drops drastically with an increase in the number of pitch angles. Also, this drop is accompanied by an increase in proportion of fields elongated along X and Y axes. However, in tilted lattice, place fields were not elongated preferentially along any axis but had an equal proportion amongst all, when the pitch angles were just 3. But as the number increased, the elongation along C axis was observed to decrease and the A, B axes shared nearly equal proportion of place fields elongated along them (Fig. 8: C1 and C2).

For aligned case, the proportion of fields oriented along Z axis dropped drastically (Fig. 8: D1) from more than expected by chance (97.5^th^ percentile) to less than expected by chance (2.5^th^ percentile) (refer Fig 6). An increase in orientation is observed along X and Y axes which plateaued as the number of angles increased. For the tilted lattice though, the expansion of pitch angle set has feeble implications as A and B axis showed a small increase which then stayed constant. C axis however saw a marginal decrease in proportion of fields oriented along it (Fig. 8: D2).

#### Helical maze and Pegboard

Hayman et al. (Hayman *et al*., 2011) carried out two experimental studies to understand the response of grid cells and place cells in 3D environments in rodents. This study used two experimental setups: a pegboard maze setup and a helical maze setup. We use the autoencoder architecture with two layers described in Methods Section (Fig. 2: B) to model the experimental results of (Hayman et al 2011).

#### Helical maze

To simulate the results, we created helical trajectories with a five-coil helix. As the results differed in upward and downward movement, the outputs of the model for both runs are considered as separate datasets.

#### Training of the model

Separate autoencoder stacks, each with two layers, are trained for upward and downward runs. The output of the PI layer is input to layer 1 of the autoencoder stack; the output of layer 1 is input to layer 2 of the autoencoder stack (Fig. 2: B). The bottleneck of the autoencoder stack comprised of layer 1 and layer 2 are probed to search for grid cells and place cell-like activity. Both the layers of the stacked autoencoders contain fifty neurons each. Similarly, a separate autoencoder stack is trained on the upward runs too. A neuron is considered to be “firing” if it crosses a specified threshold, and a red dot marks the location of its firing on a blue trajectory.

#### Results

The neurons that fire at multiple levels of the helical trajectory coils stacked at the same location on the XY plane is defined as place cells. Fig. 9 shows the top view of the helix, displaying firing rate fields and heat map of place cells. To display the firing rate of each cell over each coil, we unwound every coil into a linear strip. The firing plot is created by dividing each coil into 100 angular bins across the XY plane and creating a frequency plot of firing points for each bin.

**Figure 9:**
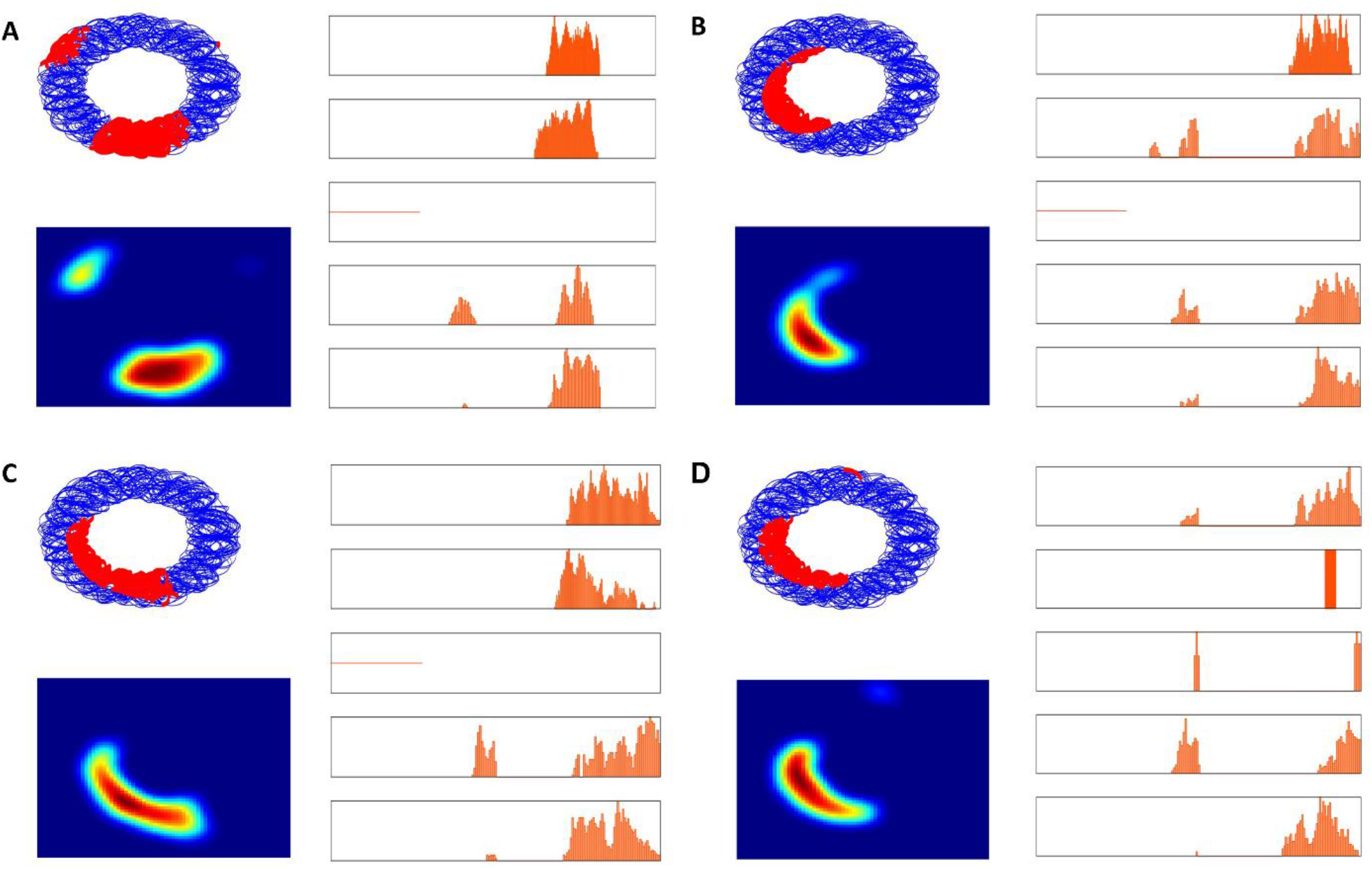
A, B, C, and D depict the activities of four sample place cells. The two columns on the left side depict the top view and firing rate map in helical maze respectively. The graphs in the right column denote the linear firing frequency on each of the 5 coils as a function of position along the track (divided into 100 bins).

The neurons which fire at more than one location on the same coil stacked over multiple coils are grid cells. Similar linear plots for each coil have also been created for grid cells (Fig. 10). We analyze both the layers of the model for grid cells and place cells. Since the firing locations for both the runs, i.e., upward run and downward run, are different, we analyze the firing pattern of these neurons in both runs (Supplementary material: section 7 Fig. S2 and S3). This observed difference in firing location in both the runs may be due to the resetting of phase at extreme points of the helical environment. Therefore, to introduce this to our model, we also reset the phase at the extreme points, i.e., the initial points for both the runs are different. For the upward run, the initial point will be the bottom-most point and for the downward run it is the topmost point.

**Figure 10:**
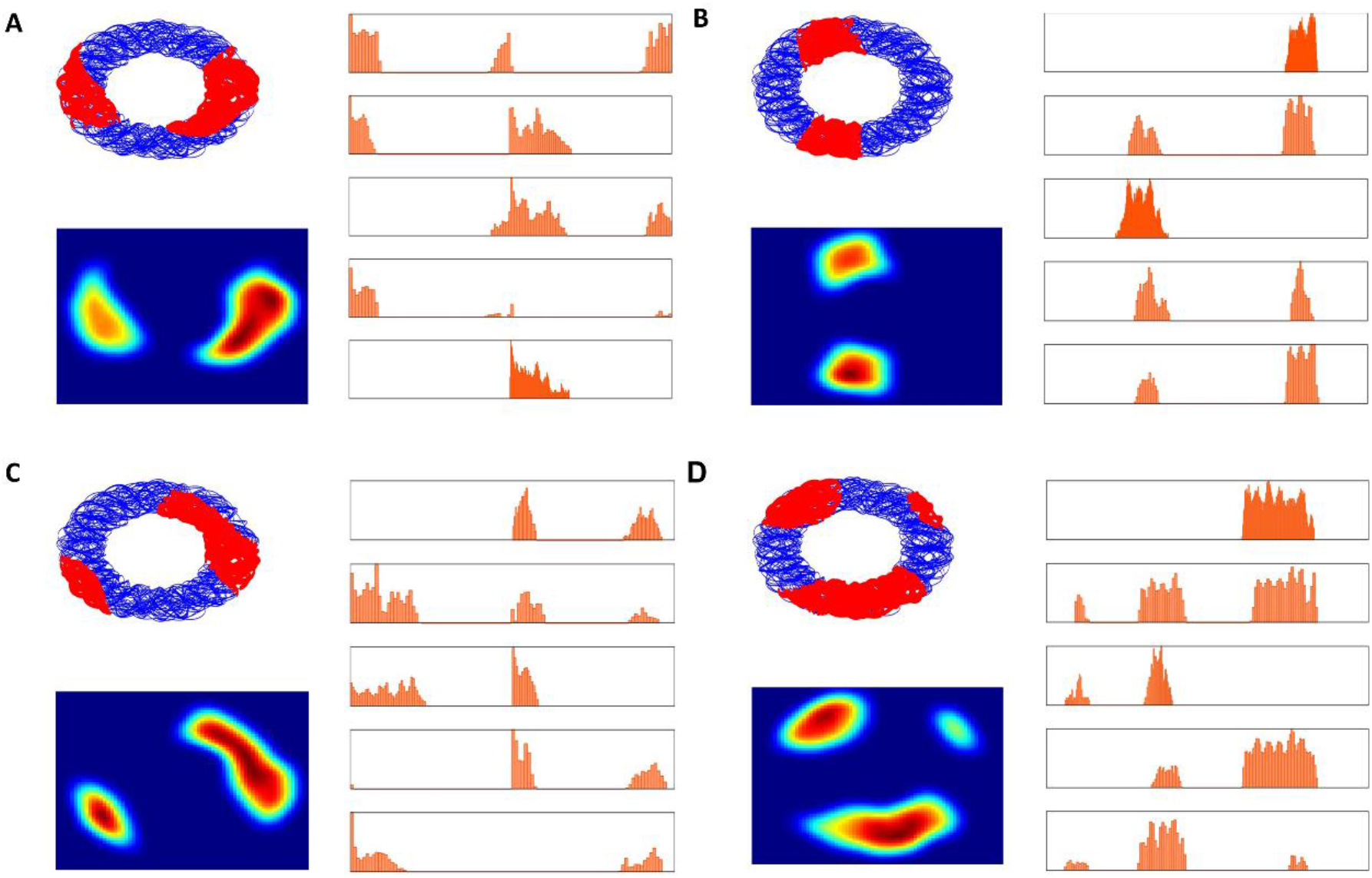
A, B, C, and D are four example grid cells. The two columns on the left side depict the top view and firing rate map in helical maze respectively. The graphs in the right column denote the linear firing frequency on each of the 5 coils as a function of position (divided into 100 bins).

In the downward run, out of the 50 neurons in layer 1 of the model, 23 grid cells and 7 place cells are observed, whereas, in layer 2, 23 grid cells and 13 place cells are observed. In the upward run, 25 grid cells and 4 place cells are observed in layer 1 of the model, whereas 23 grid cells and 18 place cells are observed in layer 2. Most of the place and grid fields are observed to span across three to five coils, and a significant number of place firing fields are observed to span four or more coils (Fig. 11).

**Figure 11:**
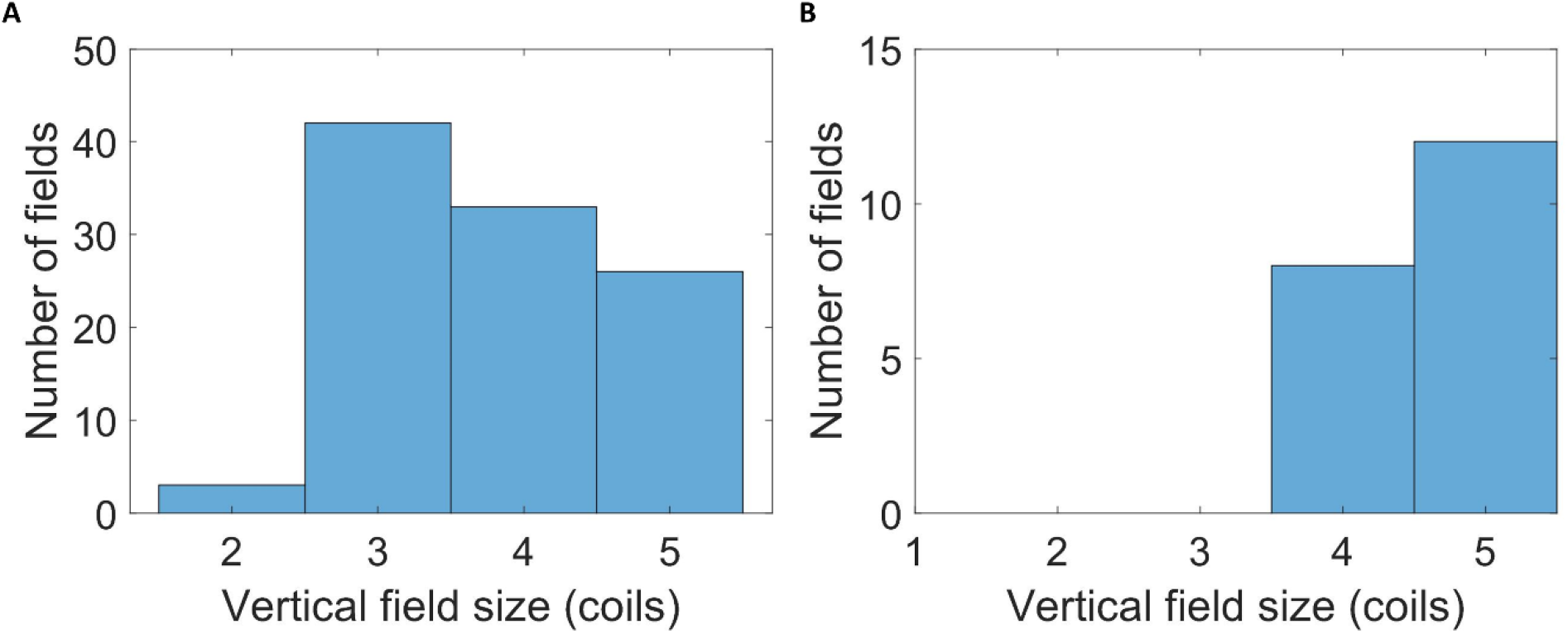
A) Vertical field size vs number of fields in observed grid cells. B) Vertical field size vs number of fields in observed place cells.

#### Pegboard Maze

We created a trajectory on a vertical maze that has pegs protruding out of a vertical board (Fig. 1: E). The agent moves around the vertical board with pegs as stepping stones.

#### Training the model

We use an autoencoder stack with two layers and analyze the results in the model’s bottleneck layers. We trained the model in a 2D square environment and tested the trained model on a 3D pegboard. The output of layer 1 of the stacked autoencoder is the input to layer 2 of the stacked autoencoder. The grid and place cells are analyzed from both the layers of the stacked autoencoder for both environments, respectively. The neurons that pass the criteria of grid cells and place cells in a 2D environment are also tested on the 3D pegboard environment.

#### Results

We observe grid cell-like and place cell-like responses in both the layers of the stacked autoencoder. Both layers of the stacked autoencoder model contain 50 neurons each. In layer 1, 23 grid cells are observed when the model is trained on a flat arena. We examined the responses of 23 grid cells on the pegboard maze. Out of the 23 neurons, 10 showed grid cell-like response on pegboard maze too. In layer 2, 8 grid cells are observed in the flat arena, and out of these 8 neurons, 5 neurons show grid-like firing in the pegboard maze too.. Grid cells on the flat arena showed hexagonal grid patterns but had elongated fields in the vertical axis that spanned across the height of the pegboard when tested on the pegboard (Fig. 12).

**Figure 12:**
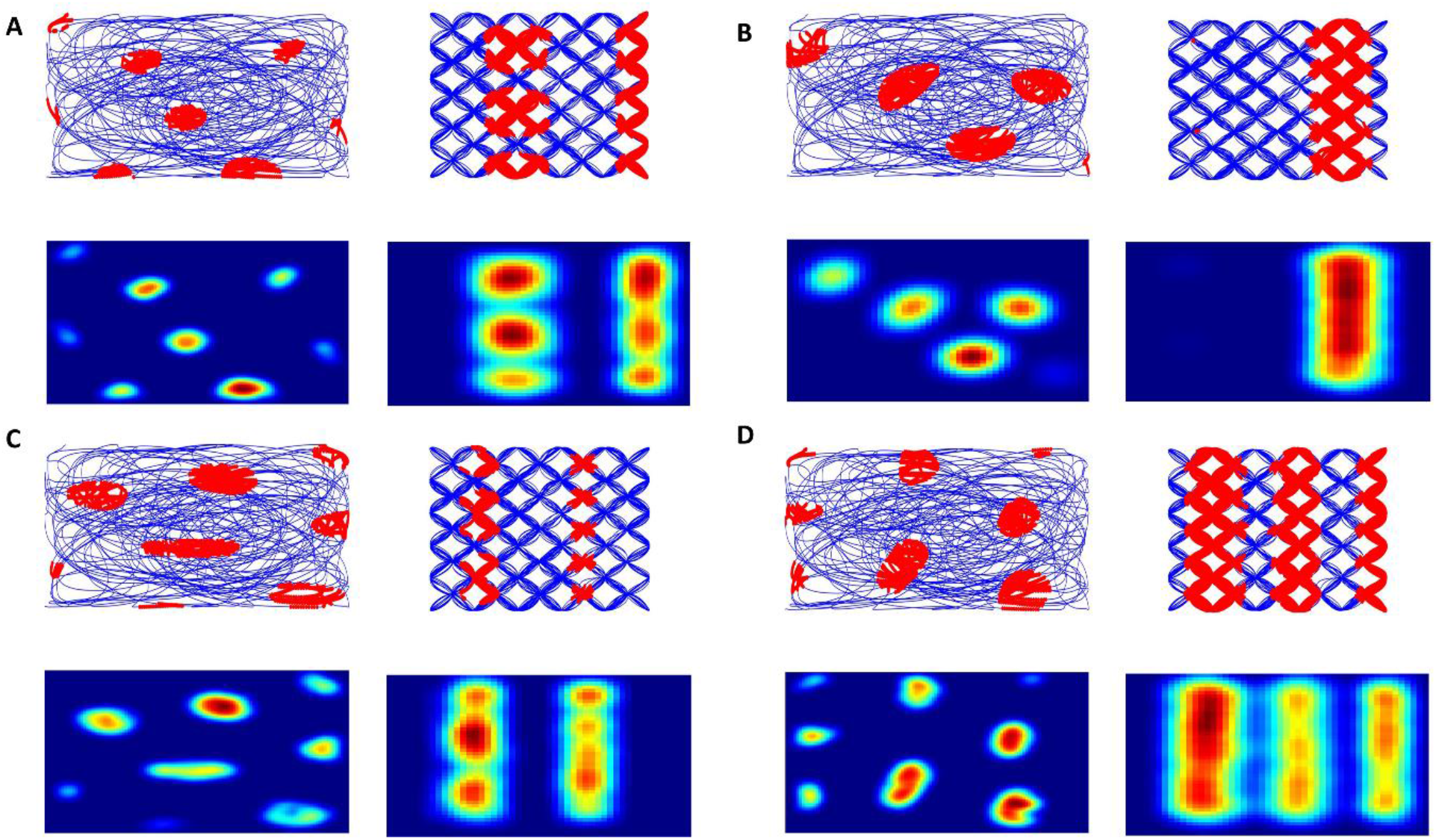
A, B, C, D are four example grid cells in the 2D environment (left) and 3D pegboard environment (right) from layer 1.

Similar to grid cells, place cell-like firing in both the layers is also observed. In layer 1, no place cell-like firing is observed when the animal is trained on a flat arena. Therefore, we did not test it on the pegboard maze. In layer 2, 12 out of 50 neurons show place cell-like firing when the agent is trained on the flat arena, and 5 out of these 12 neurons show place cell-like firing on the pegboard maze (Fig. 13).

**Figure 13:**
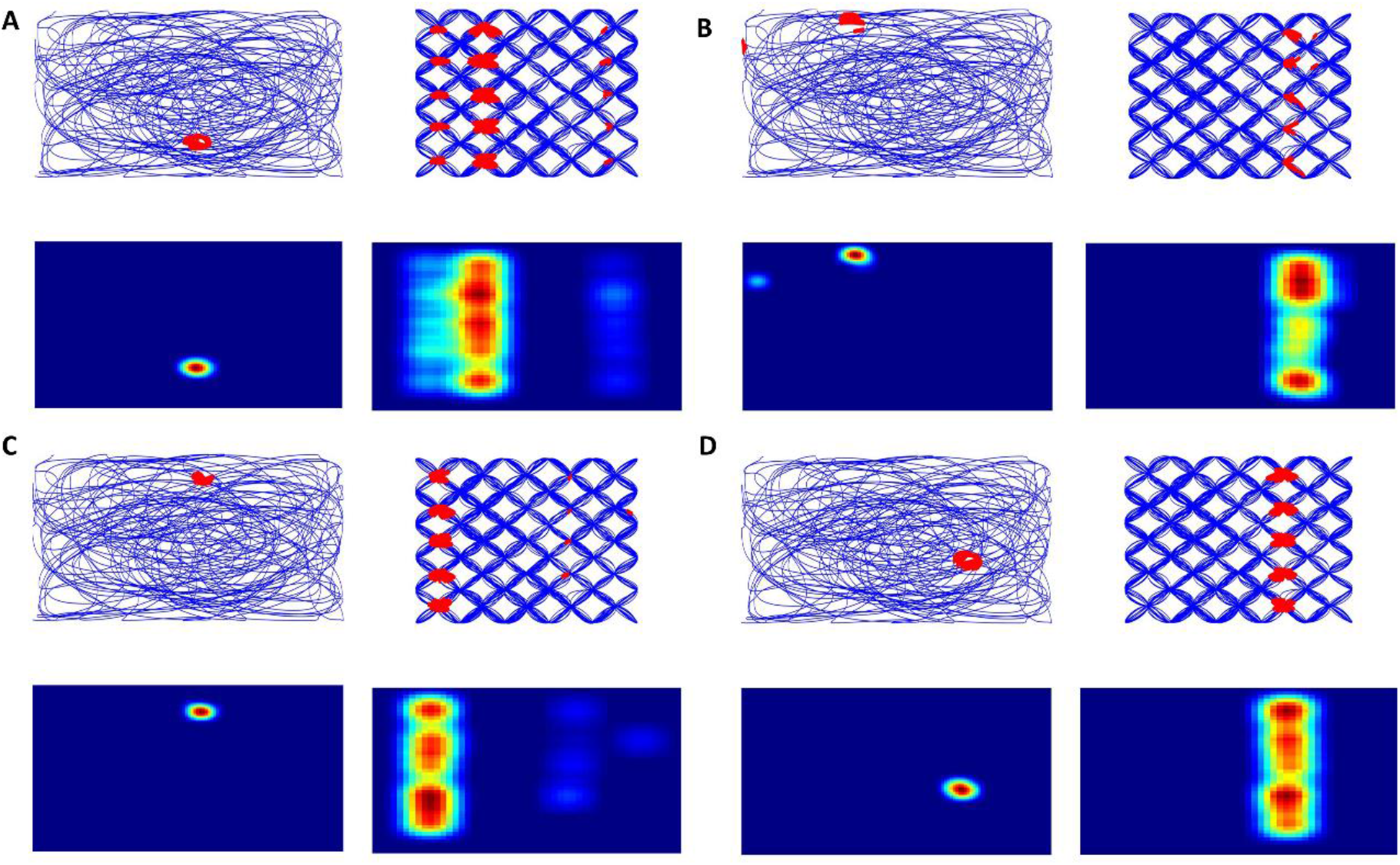
A, B, C, D are four example place cells in the 2D environment (left) and 3D pegboard environment (right) from layer 1.

#### Discussions

Computational models of the hippocampal spatial cells in 3D navigation are scarce. The few existing models are mainly abstract theoretical models of spatial cells and do not try to reproduce experimental results (Horiuchi and Moss, 2015b; Mathis, Stemmier and Herz, 2015; Stella and Trevesy, 2015). There is one study (Soman, Chakravarthy and Yartsev, 2018) that uses anti-Hebbian learning to model spatial cells like place cells, plane cells, and grid cells in case of bats navigating in 3D space. However, there is a dearth of comprehensive computational models that can model grid cells and place cells in diverse environments. Therefore, in this study, we propose an autoencoder-based deep learning approach to model place cells and grid cells in various environments. This approach has been successfully employed before to model place cells and grid cells in a 2D environment (Aziz *et al*., 2021).

First, we simulate the experimental study (Hayman *et al*., 2011) that shows rodent place cell and grid cell firing patterns when foraging on pegboard and helix. When the autoencoder model is trained on a 2D track and further tested in a 3D environment for pegboard, we observe the emergence of grid cells and place cells in autoencoder layers similar to the original experiment (Hayman et al 2011). The same was observed when we trained the model on a helical maze. The emerging spatial cells show stark spatial selectivity on horizontal planes and also the elongation observed along the vertical plane. We also observe place cells and grid cells in both the autoencoder layers. Although, it must be noted that the number of observed place cells in layer 2 of the autoencoder is significantly more than in the preceding layer 1. We can draw a speculative conclusion from the above observations that layer 2 is analogous to the CA (Cornu Ammonis) region of the hippocampus, whereas the preceding layer 1 is comparable to MEC (Medial Entorhinal Cortex).

The other study we stimulated was from (Grieves *et al*., 2020), which involved a rat’s movement along an aligned and a tilted 3D lattice maze. We simulate a trajectory that allows a rat to forage along a 3D lattice cube which can then be rotated to incorporate the settings from the original experimental study. We used this trajectory to train the autoencoder network with a single autoencoder layer. In the aforementioned simulation with pegboards and helices, we used an autoencoder with 2 layers, since we wanted to demonstrate a hierarchy in the distribution of grid cells and place cells. However, the 3D lattice experimental study only discusses place cells. Hence in this case we use only a single layer in the autoencoder network, since we probe the network only for place cells, in line with the experiment. We show that it is possible to explain all the experimental observations (emergence of place cells, distribution of place cells, elongation, orientation along axes, and preferential elongation along specific axis) using our autoencoder network model.

An important aspect of the simulations is to understand the effect of the HD on spatial cell firing in 3D. To explore that question, we extended the study by measuring the effect of HD on all the observed phenomena (specifically for orientation and elongation along a preferred axis) in lattice mazes. The selection of pitch angles for HD tuning plays a vital role in getting the desired results for specific environments. The simulations show that the HD neurons in rats are not tuned to a broad range of pitch angles compared to the azimuth angles. The results highly depend on the selection of a pitch angle set that is highly specific to how the animal explores each direction. For example, in the aligned lattice maze, the agent’s movement is restricted chiefly to 3 pitch directions. Hence, the head direction neurons are tuned to the 0, 90, and 180 degrees, respectively. In the tilted lattice maze, the animal’s movement is mainly along the now tilted axes ABC; therefore, the HD neurons are tuned to the pitch angles of 45, 90, and 135 degrees.

Similarly, this type of setting works in helical and pegboard maze too. In the helical maze, the HD neurons get tuned to mostly azimuth angles since the animal moves along the XY plane. Similarly, the neurons get tuned to those directions in a pegboard maze since the animal’s HD mainly remains vertical - upward or downward. This tuning of HD cells explains the anisotropic nature of place fields for rodents in 3D which stand in contrast to the nearly isotropic fields in bats (Hayman *et al*., 2011; Yartsev, Witter and Ulanovsky, 2011; Grieves *et al*., 2020). This trend perhaps stems from the restricted foraging of rats in 3D environments which allows the tuning of HD neurons only for accessible directions (specifically in pitch), which is marginally different from the isotropic tuning of HD in bats due to increased accessibility of directions.

From the above studies, it is evident that the limitations of rodent biology, that restrict its movements to a surface, and prevent full 3D navigation, are the primary reason for the preceding observations regarding HD tuning. Alongside, we have successfully presented the versatile nature of deep learning approach to model spatial cells in the hippocampal formation. The proposed modeling approach is quite comprehensive as it uses autoencoder layers to model place cells and grid cells in different studies and environments using a single model. This model can be further expanded to integrate other sensory inputs like vision into the model in 3D environment.

## Contribution of authors

AA contributed in modelling, coding, analysis, and manuscript writing. KL contributed in modelling coding and analysis. BKP contributed in result and statistical analysis, and manuscript writing. PS contributed in modelling, and analysis. AM contributed in ideation. VSC contributed in ideation, modelling, manuscript writing. All authors have approved and accepted the final manuscript.

