## Supplementary Material for "Modeling Hippocampal Spatial Cells in Rodents navigating in 3D environments"

1. Trajectory: Lattice maze (aligned and tilted)

Table 1: Probability of virtual agent choosing the direction of next node in aligned lattice maze

| Aligned Lattice Probabilities | |
| --- | --- |
| Preferred Movement | Probability |
| 1. Vertical movement    a. Upward movement    b. Downward movement | 20%  8%  12% |
| 2. Horizontal movement    a. Movement along with X or Y axes    b. Diagonal movement | 80%  70%  30% |
| Tilted Lattice Probabilities | |
| Preferred Movement | Probability |
| 3. Vertical movement    a. Upward movement    b. Downward movement | 33%  16.5%  16.5% |
| 4. Horizontal movement    a. Movement along with X or Y axes    b. Diagonal movement | 67%  70%  30% |

2. Place cells and firing fields

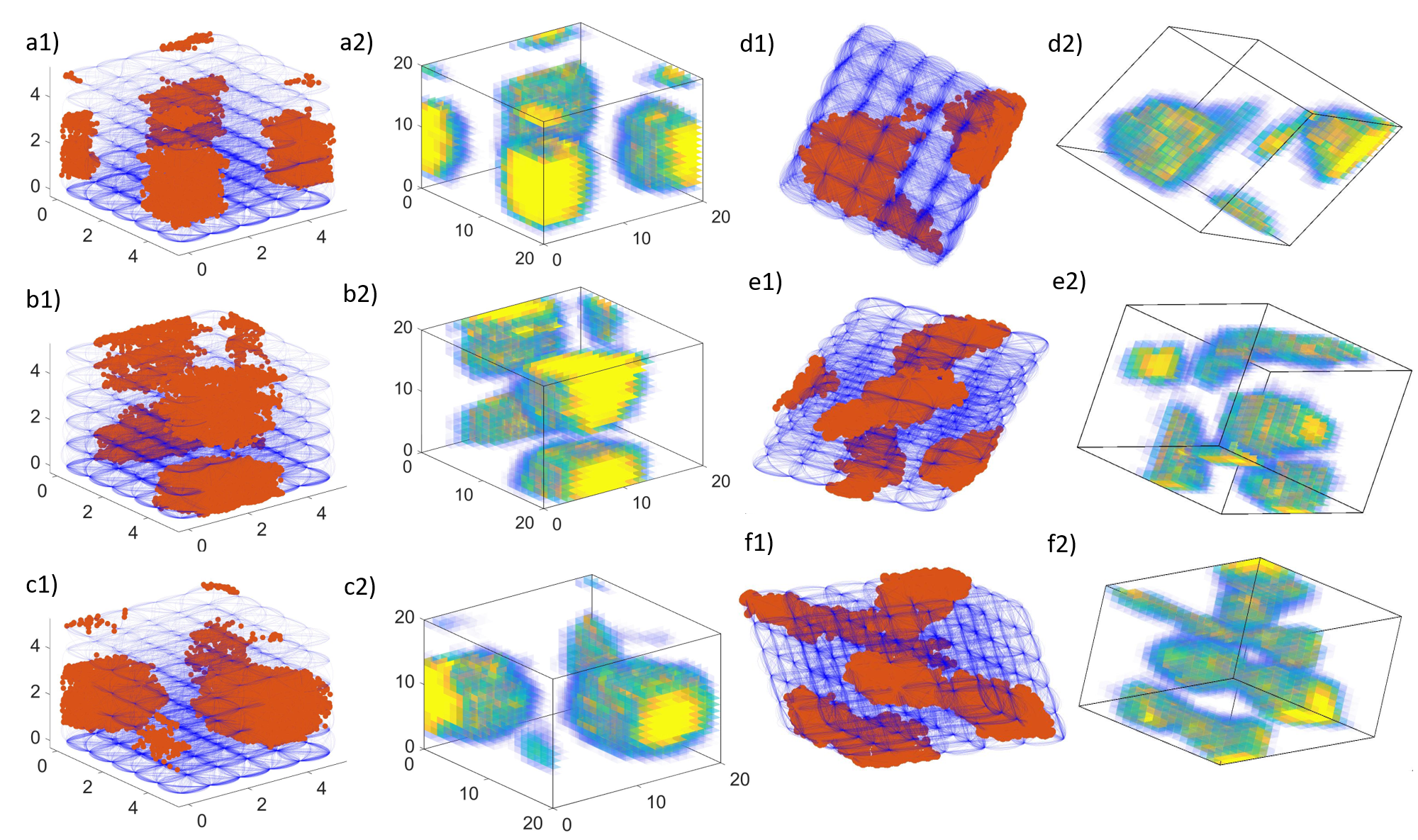

*Figure S1: a1, b1 and c1 are firing fields of three different neurons in an aligned lattice maze. a2, b2, and c2 are firing rate maps of a1, a2, and a3 respectively. d1, d2, e1, e2, f1, and f2 are similar plots for tilted lattice maze.*

3. Distribution of fields:

We extracted the centroids for all the place fields in both the lattice mazes (*regionprops3)* and calculated their medians along each axis. To test whether the fields were distributed uniformly inside the lattice volume, we created a distribution of medians for each axis. To achieve this N random points (equal to the number of fields) were generated and their median was calculated along each axis. This was performed 1000 times for each maze. If the median for a certain axis falls between 2.5^th^ and 97.5^th^ percentile of the distribution, place fields were considered to be uniformly distributed along that axis (Fig 4 a3, b3).

4. Field Elongation:

To compare the elongation of the place field for both mazes we calculated the elongation indices for all place fields. The former was achieved by fitting an ellipsoid around it and calculating the principal axes components for each place field as inspired from (supplementary material (Grieves et al. 2020)). These principal axes were defined as the major axes of the ellipsoid. Further, an elongation index for each place field is defined as

$$\boldsymbol{Field Elongation Index=}\frac{\boldsymbol{2}\boldsymbol{P}_{\boldsymbol{1}}}{\boldsymbol{P}_{\boldsymbol{2}}\boldsymbol{+}\boldsymbol{P}_{\boldsymbol{3}}}$$

Where, *P_1_*, *P_2_* and *P_3_* are principal axes in descending order. The above elongation index indicated the spherical nature of a field, with elongation index of 1 indicating spherical field. A majority of fields were elongated in both aligned (Mean: 2.00, Std. Error: 0.54) and titled (Mean:1.94, Std. Error: 0.61) lattice.

To test the deviation of elongation index from a distribution that would be expected by chance we did the analysis illustrated in the experimental result (Grieves et al. 2020). We created a sphere with the center being the centroid of each place field. The diameter of the sphere is given as

$$\boldsymbol{Equivalent diameter=6}\left( \frac{\boldsymbol{v}_{\boldsymbol{f}}}{\boldsymbol{п}} \right)^{\frac{\boldsymbol{1}}{\boldsymbol{3}}}$$

Where, the trajectories that would lie inside the sphere were screened. A set of normally distributed random points equal to the number of spikes of the place field were generated inside the sphere. The mean of the distribution was the centroid and standard deviation of 1.4 times the equivalent radius. Further each of these points were assigned to a closest point on trajectories screened inside the sphere. This created a normal distribution of spikes inside the sphere. Place fields were identified from this random shuffling of spikes and field elongations were calculated. The shuffling was performed 100 times for each place field. From probability distribution for each place field we observed whether the elongation index for each field was greater or equal to expected by chance. If the actual elongation fell under the 95th percentile of the distribution, it was considered spherical otherwise non-spherical.

5. Orientation:

To quantify the orientation of place fields, we test whether they were oriented along axes of interest and to differentiate the axes with respect to orientation, we performed the analyses and tests presented in the experimental study (Grieves et al. 2020). The primary principal axis of each place field, also considered as the major axis for the ellipsoid (*field elongation),* was extracted using PCA (*regionprops3*). These vectors were then projected onto a sphere with unit radius on diametrically opposite ends. To test for orientation along a specific axis, we screened the points that fell within those surface regions of the sphere that formed a ~60^o^ cone with apex at origin and axis of interest as cone’ axis. All the points from both directions for each axis were summed and presented as a ratio of total points on the sphere.

To determine whether the axes differed from each other, we calculated 95% confidence intervals for each axis. We sampled a set of orientations for randomly chosen place fields with replacement, such that the set is equal to the total number of fields. This was performed 1000 times for each maze, and ratios for each axis were recalculated. If the proportion of points parallel to a certain axis falls inside the error bars of other axes, they are not considered significantly different. The error bars in (Fig 6 a3, b3) show the 2.5^th^ and 97.5^th^ percentile for the shuffled counting.

To check whether this ratio for a certain axis was observed more than expected by chance we generated 1000 random points on the surface of the sphere. We calculated the aforementioned proportions for all the axes. We did this 1000 times and calculated the chance as 2.5^th^ and 97.5^th^ percentile of the distribution. The proportion for the axis that exceeded the upper threshold, was considered to have shown significant orientation parallel to a certain axis.

Similar to the experiment study, to check if fields were parallel to XYZ axes in aligned case and to ABC axes in the tilted case, we defined the axis ratio.

Axis Ratio = XYZ / ABC

This ratio was calculated by generating 1000 random points on the surface of a sphere 1000 times. The proportions for each axis were calculated for each shuffle and the distribution for the above ratio was plotted. If the actual ratio deviated from the 1^st^ or 99^th^ percentile, it was considered a significant deviation from the no axis bias condition.

6. Binary Morphology:

The firing rate maps of the place cells are created by first thresholding them at 10% of their maximum value, and a morphological erosion process was performed on the 3D binary volumetric image using structuring element vectors of varying lengths along each cartesian axis. For each erosion operation, the linear sum of the remaining voxels as a measure of the map’s connectivity along that dimension is first calculated and is then expressed as the proportion of all remaining voxels for that element length.

Using this approach, we observed for the aligned lattice that the connectivity of voxel was identical (nearly 33% for all axes) to a specified length of structuring element vector. It started diverging for longer elements, with a larger proportion of fields showing connectivity in the Z-axis compared to others. A repeated measures (within-subjects) ANOVA was run with axes as an independent variable and the proportion of voxels (connectivity) as a dependent variable. The results suggest a significant difference of voxel proportion between all three axes (effect of axes: F(2, 587) = 80.35, p < 0.0001, $\boldsymbol{\eta}_{\boldsymbol{p}}^{\boldsymbol{2}}$= 0.169, effect of axis and length of structuring element: F(27, 587) = 2.94, p < 0.0001, $\boldsymbol{\eta}_{\boldsymbol{p}}^{\boldsymbol{2}}$= 0.188). Post hoc tests (Bonferroni correction for pairwise Comparison) for the same revealed difference between voxel proportions for the X-axis (Mean: 0.275, Std. Error: 0.008) and Y-axis (Mean: 0.272, Std. Error: 0.008) as insignificant (did not differ), but significant difference of connectivity with the Z-axis (Mean: 0.453, Std. Error: 0.012). Pairwise comparisons show (X vs Y: p > 0.99, X vs Z and Y vs Z: p < 0.0001).

Similarly, for the tilted lattice case, the proportion of voxel connectivity stayed equal (nearly 33%) for all the axes. A slight divergence is observed for longer structural elements but is less significant as compared to aligned lattice cases. The similar repeated measures ANOVA test was run with axes as an independent variable and proportion of voxels (connectivity) as a dependent variable. The result suggested a significant difference for voxel proportion between all three axes (effect of axes: F(2, 1838) = 5.65, p < 0.005, $\boldsymbol{\eta}_{\boldsymbol{p}}^{\boldsymbol{2}}$ = 0.006) but with comparatively smaller effect size. For interaction between axis and structure length, the result showed no significant difference (F(36, 1838) = 0.25, p > 0.99, $\boldsymbol{\eta}_{\boldsymbol{p}}^{\boldsymbol{2}}$ = 0.005). Post hoc tests (Bonferroni correction for pairwise Comparison) revealed differences between voxel proportions for the A-axis (Mean: 0.352, Std. Error: 0.008) and B-axis (Mean: 0.339, Std. Error: 0.008) as insignificant (did not differ), and comparatively significant difference of connectivity with C-axis (Mean: 0.309, Std. error: 0.008). Pairwise comparisons show (A vs B: p > 0.99, B vs C and A vs C: p < 0.005).

7. Grid cells and Place cells for helical trajectory:

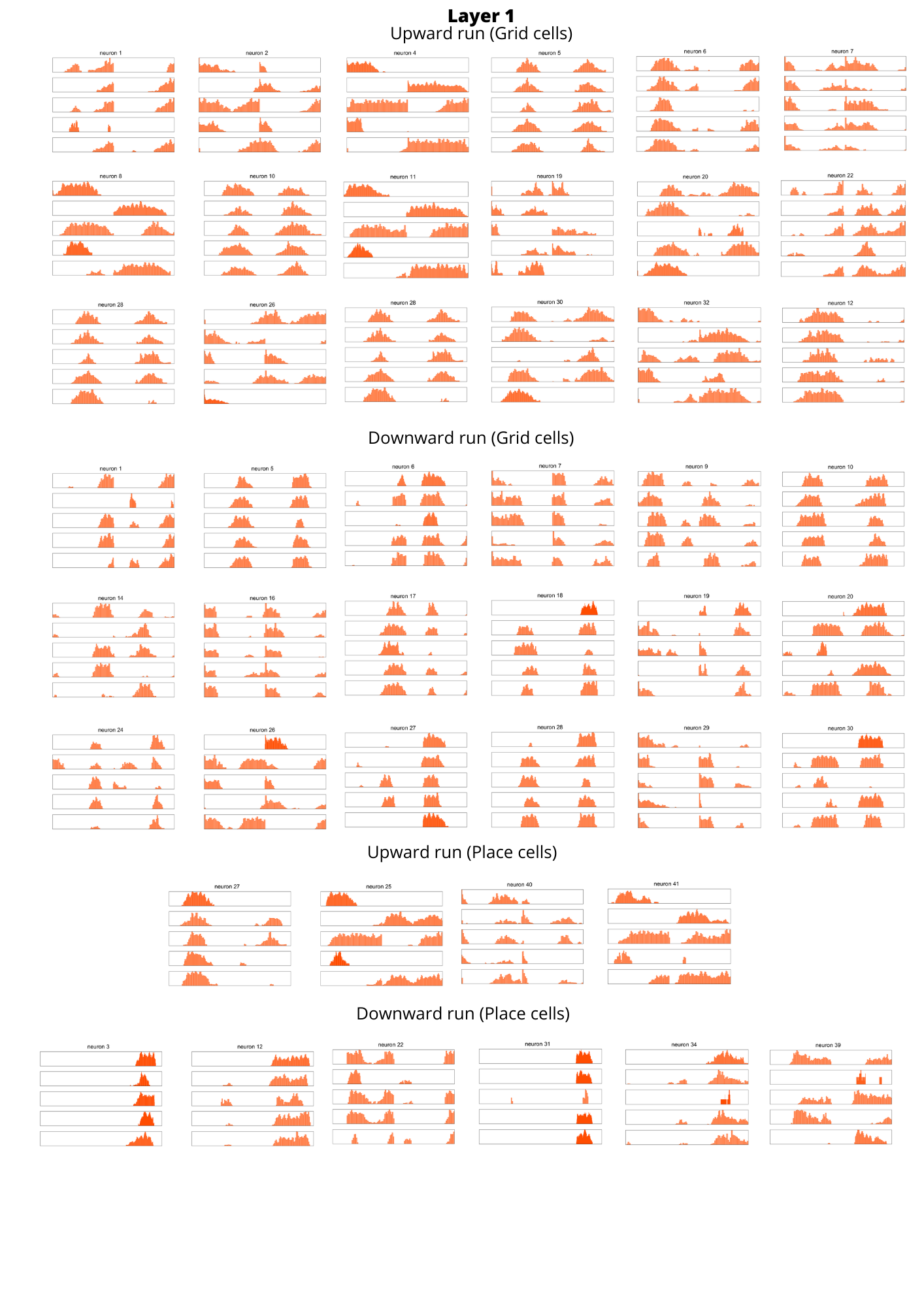

*Figure S2: Linearized grid cells from layer 1 for each coil recorded during upward and downward runs(first six rows). Linearized place cells from layer 1 for each coil recorded during upward and downward runs (last two rows)..*

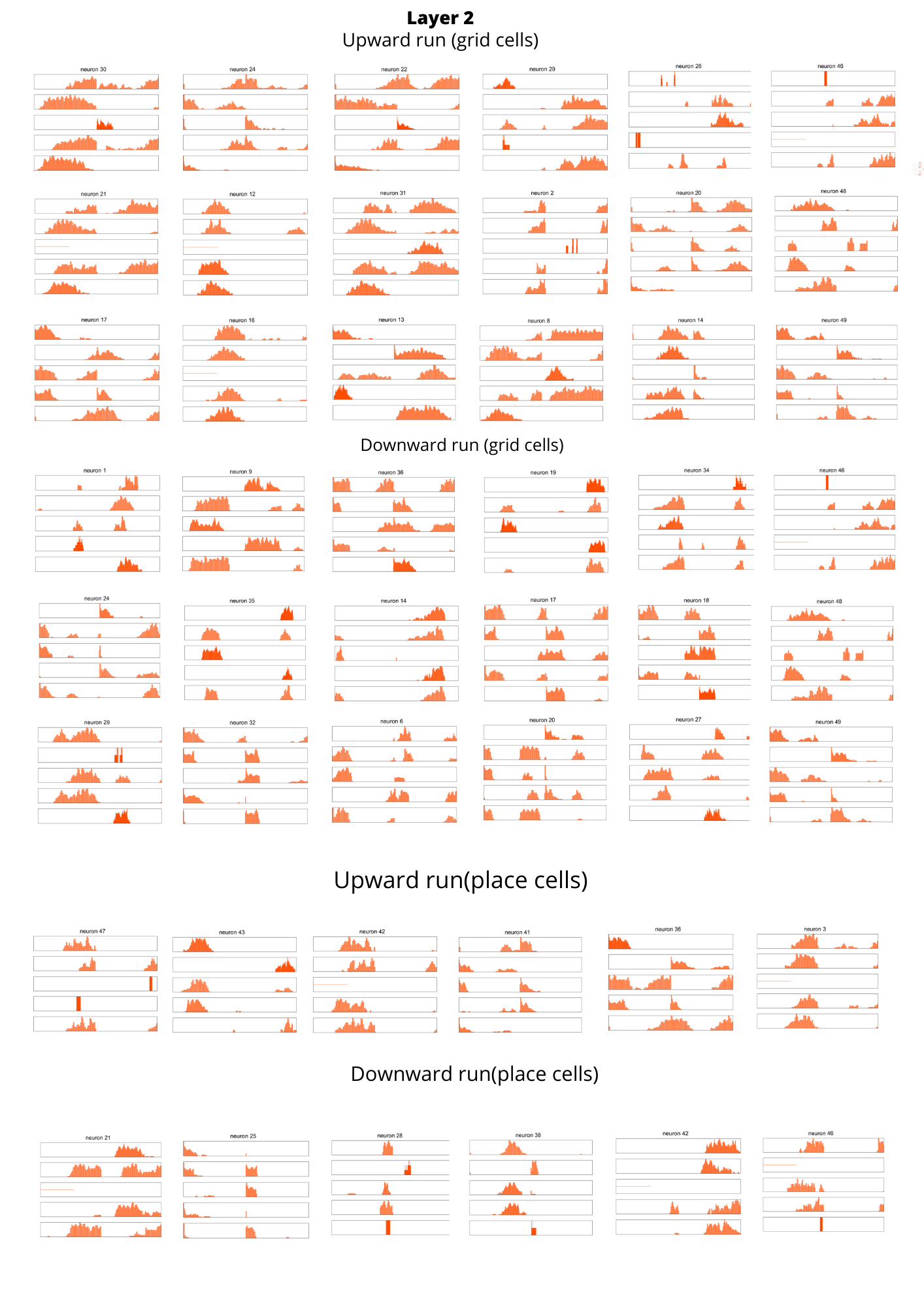

*Figure S3: Linearized grid cells from layer 2 for each coil recorded during upward and downward runs(first six rows). Linearized place cells from layer 2 for each coil recorded during upward and downward runs (last two rows).*

.
